# Evolutionary rescue from antagonistic invaders: Birth-limiting competition among residents can aid rescue

**DOI:** 10.64898/2026.01.14.699172

**Authors:** Alexander Longcamp

## Abstract

From invasive predators displacing wildlife to medically deployed bacteriophages eliminating pathogens, biological invasions by antagonists are a fundamental driver of extinction, with implications for both conservation and medicine. Resident populations facing such invasions can sometimes avoid extinction through adaptation, a process known as evolutionary rescue. Here, an analytical approximation is derived for the probability of evolutionary rescue from antagonistic invaders, focusing on how rescue depends on a key antagonist trait—victim specificity. Rescue is shown to be most likely either when invaders are slow-spreading, resident-dependent specialists whose failure to encounter suitable victims buys time for mutation, or when invaders are fast-spreading generalists that rapidly relieve standing genetic variation of intraspecific competition. The central result is that, whether invaders are generalists or specialists, a lower resident birth rate can facilitate rescue relative to a higher one by diminishing the effect of demographic stochasticity on mutant establishment. If the lower birth rate is density-independent, it primarily promotes rescue from generalists, whereas if it arises from birth-limiting competition it can also appreciably aid rescue from specialists. Overall, the results suggest that invader victim specificity can reshape evolutionary rescue by dynamically coupling the rate of resident decline to the growth rate of the stressor driving the decline, with resident life history mediating the outcome.

## 1 Introduction

Biological invasions by antagonists—whether predators, parasites, or other exploitative organisms—are a fundamental driver of extinction that can harm or protect native communities depending on the context. Invasive predators, for example, are a leading cause of the species extinction crisis, accounting for over half of all documented contemporary bird, mammal, and reptile extinctions (Doherty et al., 2016). Conversely, humans sometimes introduce antagonistic biocontrol agents that displace invasive species, disease vectors, or pests (De Clercq et al., 2011). This dual impact of antagonist invasion is also evident at the cellular scale: pathogens continually invade healthy cell populations, while medical practitioners sometimes deploy bacteriophages that infect and eliminate bacterial pathogens (Sulakvelidze et al., 2001). Understanding how and when populations might be able to withstand such invasion can thus inform strategies in both conservation and medicine.

Although antagonist invasion typically occurs on short, ecological timescales, there is evidence that populations can evolve to withstand it (Marshall & Fenner, 1958; Strauss et al., 2017; Zuk et al., 2006). For example, field crickets on the Hawaiian Island of Kauai were approaching extinction due to the invasion of a deadly parasitoid fly until a mutation prevented the male crickets from chirping, which in turn prevented the males from attracting the parasitoid (Tinghitella, 2008; Zuk et al., 2006). The Australian population of the invasive European rabbit was declining due to the controlled introduction of the Myxoma virus until the rabbits evolved greater innate immunity to the virus (Marshall & Fenner, 1958). Furthermore, laboratory populations of plankton rebounded from near-extinction by evolving to overcome the dual pressures of an invading fungal parasite and competition with a less susceptible host (Strauss et al., 2017).

This process, where a population avoids extinction through adaptation, is known as evolutionary rescue (ER; Gomulkiewicz & Holt, 1995). In its simplest form, ER entails a resident population declining toward extinction until an adaptive mutant lineage—one from either standing (genetic) variation or *de novo* mutation—establishes by overcoming demographic stochasticity (Orr & Unckless, 2008). ER theory involving antagonistic interactions often only implicitly includes biological invasion as the cause of resident decline, focusing instead on how ER dynamics are influenced by interactions between members of a focal resident species (hereafter, residents) and those of another established species (Osmond et al., 2017; Shang et al., 2024; Vanselow et al., 2022; van Velzen, 2023; Yamamichi & Miner, 2015). However, a growing theoretical literature has begun to explicitly address antagonist-driven ER, with studies considering invaders in the form of predators (DeLong & Belmaker, 2019), carriers of a pathogen (Golas et al., 2021; Jiao et al., 2020), and exploiters of resident-constructed habitats (Longcamp & Draghi, 2025).

Despite these theoretical advances, few ER models have accounted for a fundamental feature of antagonistic invaders: the degree to which their proliferation depends on a specific host, prey, or other victim species—hereafter, victim specificity. Some invaders are generalists that can proliferate independently of residents; for example, the brood parasitic shiny cowbird, which recently invaded Puerto Rico, parasitizes a wide range of Puerto Rican bird species (Cruz et al., 1985). Other invaders are specialists that depend on residents, as with the emerald ash borer—an invasive beetle that feeds almost exclusively on ash trees in its introduced range (Sun et al., 2024). Both generalists and specialists frequently succeed in invading, often because they lack natural enemies (Roy et al., 2011) or because local victims are initially unadapted to their presence (Rebek et al., 2008). Relative to generalist invaders, however, specialists tend to displace a given population more slowly due to their narrower range of suitable victims (Stiling & Cornelissen, 2005). This limitation is well documented in biological control, where specialists are often considered less effective but safer agents than generalists because they are less capable of exploiting non-target species (Stiling & Cornelissen, 2005; Van Lenteren et al., 2003). The same limitation has been illustrated in two-species microbial predator-prey experiments: when prey densities were low due to strong intraspecific competition, predators had a lower chance of encountering prey and consequently drove them extinct more slowly (Holyoak, 2000; Luckinbill, 1974).

Here, an analytical approximation is derived for the probability of ER, *P*_*ER*_, from antagonistic invaders that can be generalists or specialists. The study begins with a formulation of an eco-evolutionary model of resident-invader dynamics that is then modified for ER via an invader-resistant mutation (Figure 1). In the model, ancestral residents (hereafter, wild types) are harmed by resident-invader interactions and intraspecific competition, both of which can range from interactions that limit successful births (i.e., limit fecundity or offspring survival; e.g., brood parasitism, predation on offspring) to those that promote adult mortality (e.g., predation, aggression). For simplicity, invaders are assumed to have the same effect on wild type fitness as intraspecific competition, meaning that they displace wild types through high densities (e.g., those achieved by emerald ash borers; McCullough, 2020) rather than a strong per-capita effect on wild type fitness. Furthermore, following previous theory, invaders are assumed to have a negligible death rate (e.g., one stemming from enemy release) such that, without ER, the resident population always goes extinct in the presence of invaders (Longcamp & Draghi, 2025). Given these assumptions, *P*_*ER*_ is approxi-mated and analyzed, first with consideration of how it changes with the invader growth rate. Results show that *P*_*ER*_ is highest when invaders are either slowly invading specialists or rapidly proliferating generalists (Figure 2): slow invasion provides time for *de novo* mutations to appear, while fast invasion rapidly releases standing variants from competition with wild types. Next, *P*_*ER*_ is analyzed as a function of the resident birth and death rates. The main finding of the study is that a reduction in the resident birth rate—but not a boost in the death rate—can enhance *P*_*ER*_ (Figure 3) by lessening the effect of demographic stochasticity on mutant establishment. If the reduced birth rate is density-independent, it primarily increases *P*_*ER*_ when invaders are generalists (Figure 3), whereas if it stems from birth-limiting competition it can appreciably raise *P*_*ER*_ under both generalist and specialist invasion (Figure 4). Even when competition-induced, however, the lower birth rate is only beneficial if mutants are sufficiently sensitive to death-promoting competition (Figure 5).

**Figure 1.**
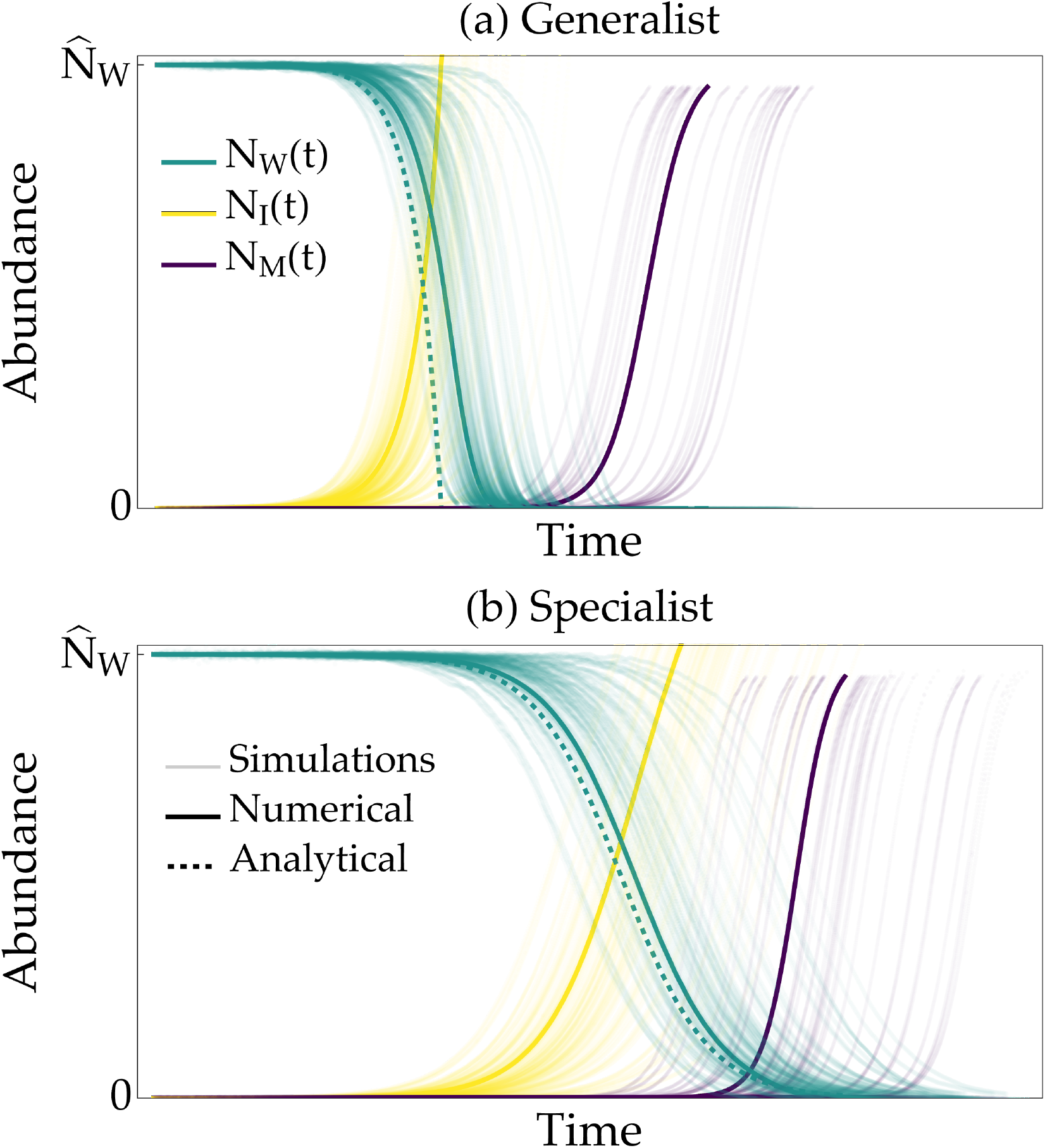
Stochastic and expected dynamics of evolutionary rescue. Panel (a) shows the case of generalist invaders with a per-capita growth rate of *r*_*I*_ (*t*) = Φ_*G*_, while panel (b) shows the case of specialists with *r*_*I*_ (*t*) = *n*_*W*_ (*t*)Φ_*S*_ . In both cases, wild types (*N*_*W*_ (*t*)) start at their expected equilibrium abundance, 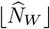, and decline toward extinction as invaders (*N*_*I*_ (*t*)) exploit them. This decline is slower with specialists because their growth rate decreases with wild type density. Before wild types go extinct, they may give rise to invader-resistant mutants (*N*_*M*_ (*t*)), either via standing variation or *de novo* mutation. Rescue occurs if 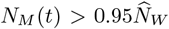 . Thin transparent curves: 100 replicate simulations of the algorithm described in Section 3.1.2, except with each simulation proceeding until *N*_*M*_ (*t*) = 0 and 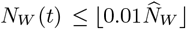 or 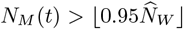; thick solid curves: numerical solution to Eq.(3), calculated with an invader carrying capacity of 100*K* to avoid integer overflow; thick dashed curves: Eq. (9). Parameter values are Φ_*G*_ = Φ_*S*_ = 0.75, *b* = 1, *d* = 0.03, *δ*_*b*_ = *δ*_*d*_ = 1, *K* = 10^5^, and *U* = 0.1.

**Figure 2.**
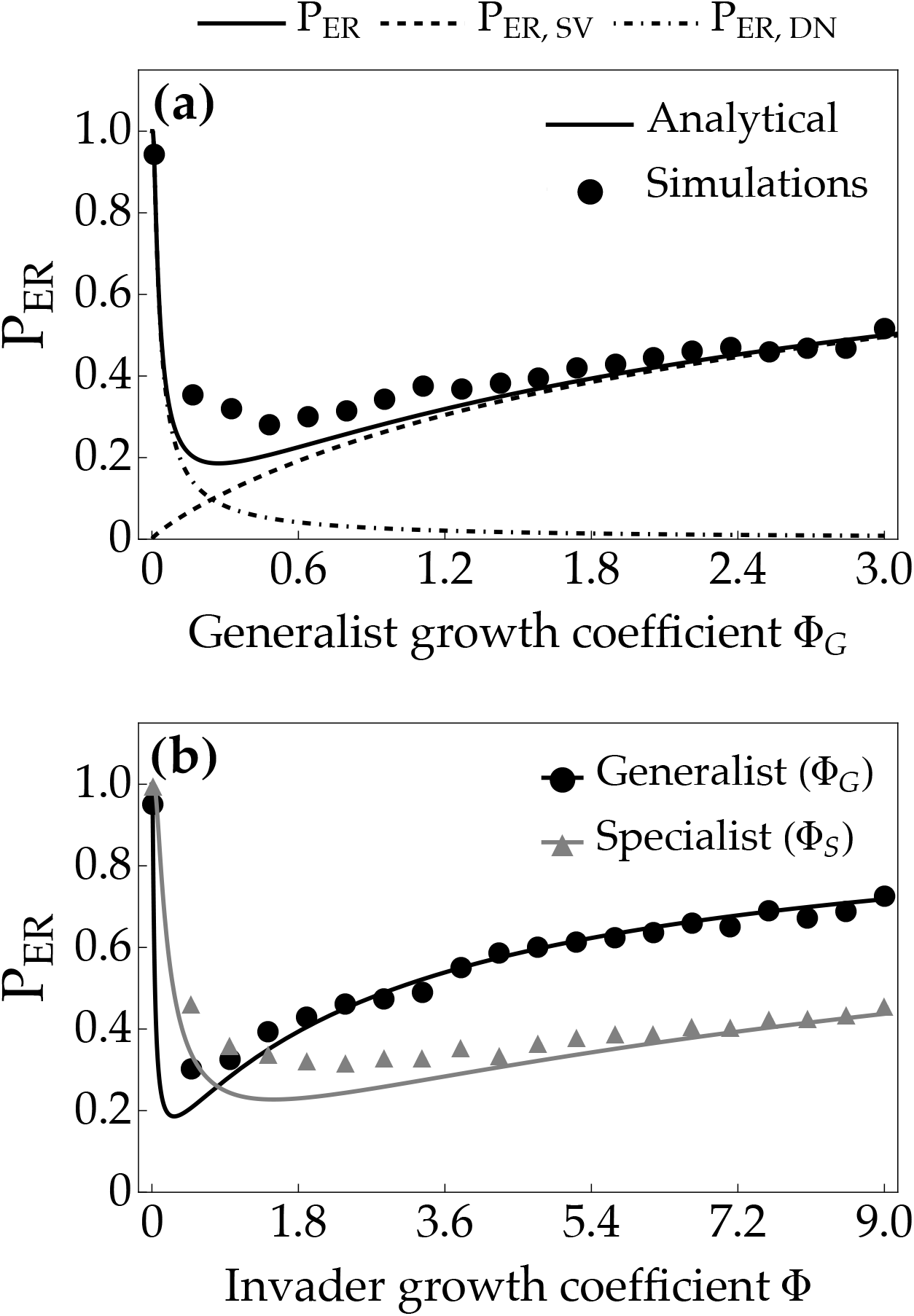
The probability of evolutionary rescue, *P*_*ER*_; the probability of rescue via standing variation, *P*_*ER*_,_*SV*_; and the probability of rescue via *de novo* mutation, *P*_*ER,DN*_, as functions of the generalist invader growth coefficient, Φ_*G*_, and the specialist invader growth coefficient, Φ_*S*_ . (a) Slow invasion favors rescue via *de novo* mutation because wild types decline slowly, providing more time for mutation; fast invasion favors rescue via standing variation because mutants are rapidly released from competition with wild types. (b) For this reason, *P*_*ER*_ is highest when invaders are either slowly invading specialists that often fail to encounter their wild type victims or rapidly invading generalists that can proliferate independently of wild types. Markers: proportion of 2 × 10^3^ replicate simulations that ended in mutant survival; dashed curve: Eq. (10); dot-dashed curve: Eq. (11); solid curves: analytical approximation of *P*_*ER*_ using Eqs. (10) and (11). Unless stated otherwise, parameter values are *b* = 1, *d* = 0.03, *δ*_*b*_ = 3, *δ*_*d*_ = 1, *K* = 10^5^, Φ_*G*_ = Φ_*S*_ = 0, and *U* = 0.1.

**Figure 3.**
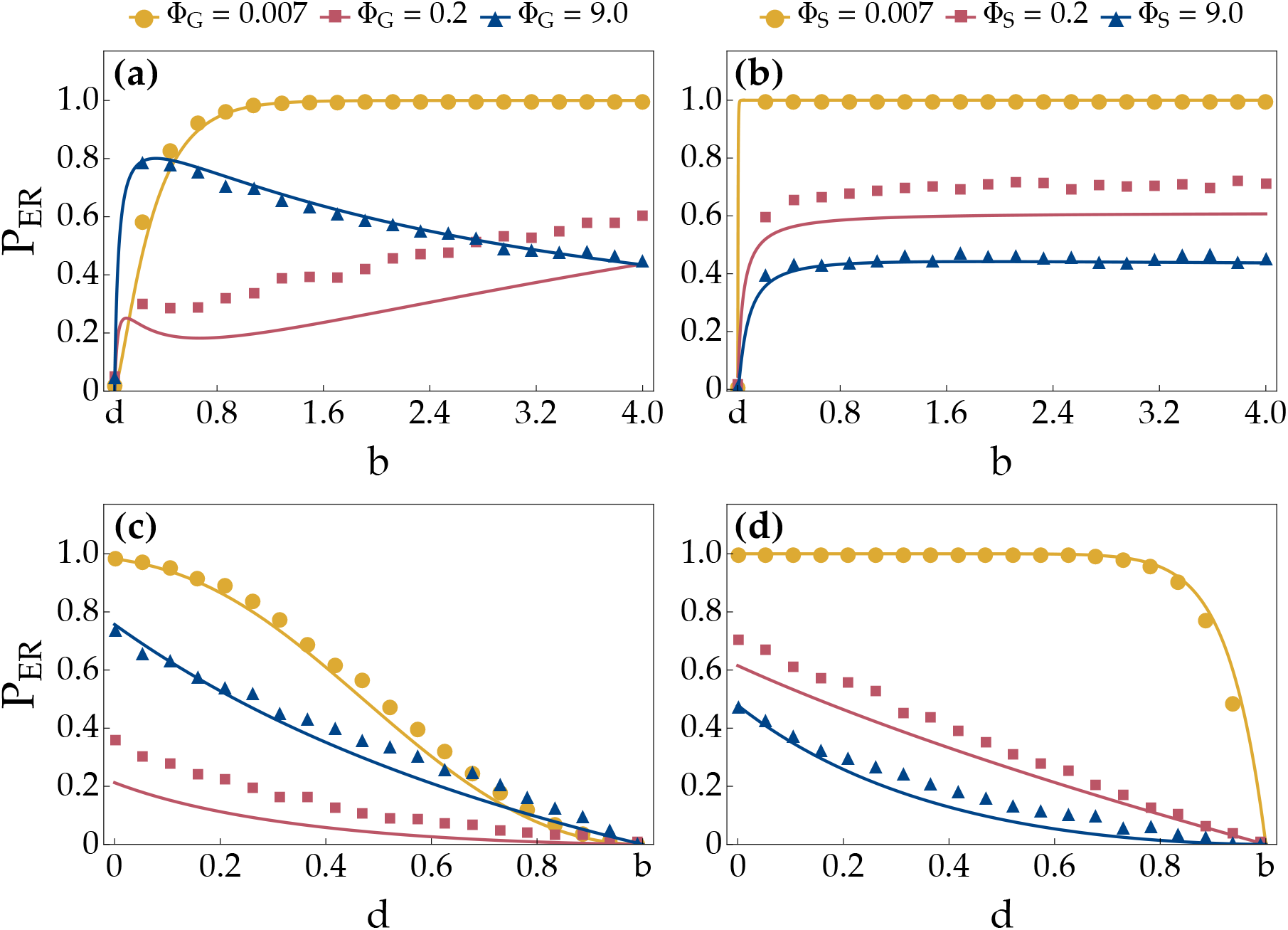
The probability of evolutionary rescue, *P*_*ER*_, as a function of the (a–b) resident density-independent birth rate, *b*, and (c–d) death rate, *d*, given different generalist invader growth coefficients, Φ_*G*_, and specialist coefficients, Φ_*S*_ . (a) For slowly invading generalists, *P*_*ER*_ increases with *b* due to greater mutational input, whereas for rapidly invading generalists, reducing *b* can increase *P*_*ER*_ by weakening the effect of wild type density on variance in mutant fitness. (b) For specialist invaders, the latter benefit does not appreciably enhance *P*_*ER*_, because fewer wild types means slower invasion, which induces a higher wild type impact on mutant fitness that offsets the benefit of lower mutant variance. (c–d) Increasing *d* never enhances *P*_*ER*_, because a higher rate of random mutant deaths means greater mutant variance. Markers: proportion of 2 × 10^3^ replicate simulations that ended in mutant survival; solid curves: analytical approximation of *P*_*ER*_ using Eqs. (10) and (11). Unless stated otherwise, parameter values are *b* = 1, *d* = 0.03, *δ*_*b*_ = 3, *δ*_*d*_ = 1, *K* = 10^5^, Φ_*G*_ = Φ_*S*_ = 0, and *U* = 0.1.

**Figure 4.**
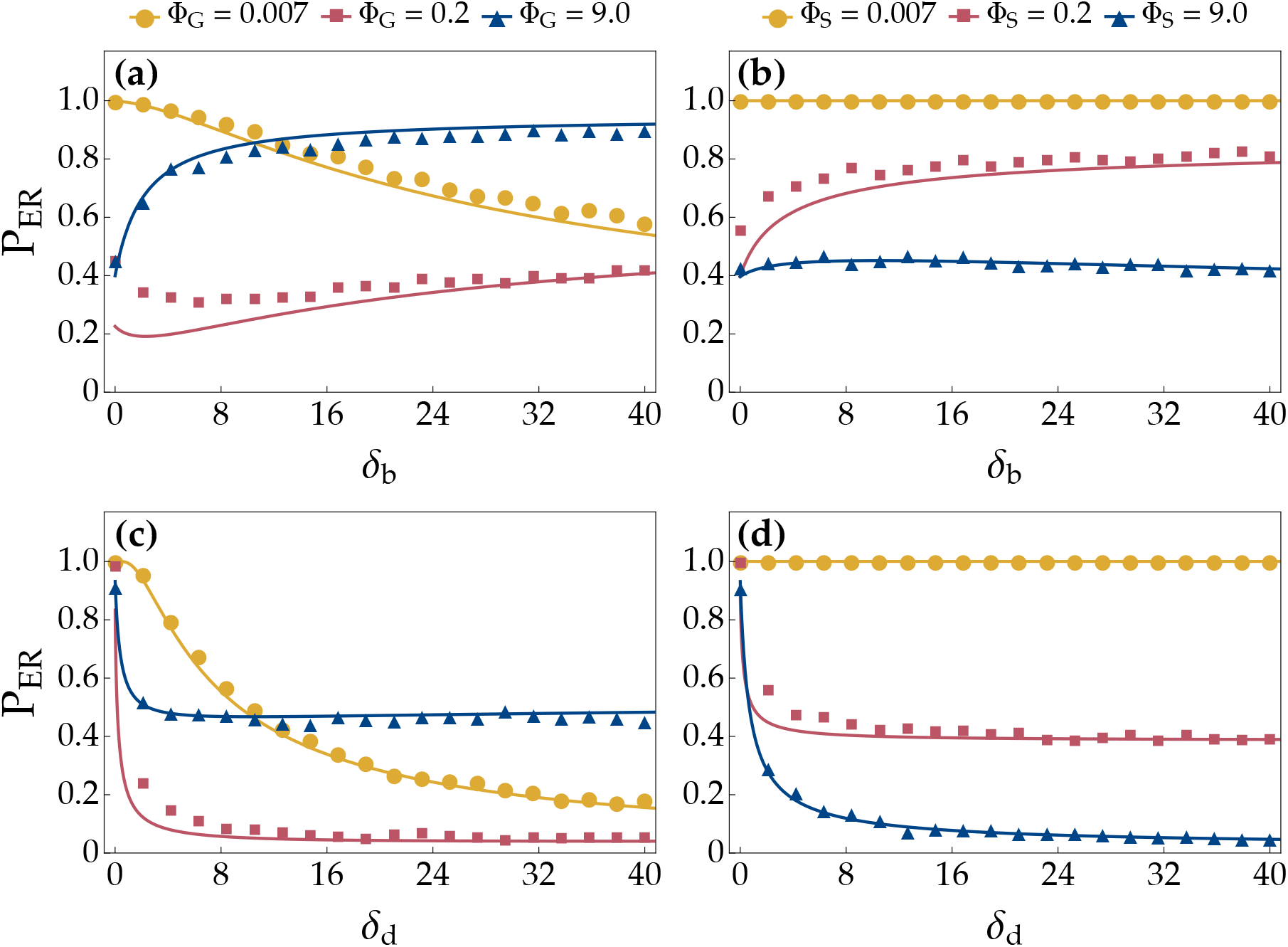
The probability of evolutionary rescue, *P*_*ER*_, as a function of (a–b) resident sensitivity to birth-limiting interactions, *δ*_*b*_, and (c–d) death-promoting interactions, *δ*_*d*_, given different generalist invader growth coefficients, Φ_*G*_, and specialist coefficients, Φ_*S*_. (a) Increasing *δ*_*b*_ can enhance *P*_*ER*_ by lowering the effect of wild type density on variance in mutant fitness. This can occur whether invaders are rapidly invader generalists or (b) slowly invading specialists. In the latter case, the mutational cost of fewer wild types is offset by a reduction in invasion speed, while the mutant fitness cost of slower invasion is weakened by the decay in the intensity of intraspecific competition as wild types decline, allowing a higher *δ*_*b*_ to facilitate rescue via *de novo* mutation. (c–d) Because a higher rate of random mutant deaths means greater mutant variance, increasing *δ*_*d*_ decrease *P*_*ER*_. Markers: proportion of 2 × 10^3^ replicate simulations that ended in mutant survival; solid curves: analytical approximation of *P*_*ER*_ using Eqs. (10) and (11). Unless stated otherwise, parameter values are *b* = 1, *d* = 0.03, *δ*_*b*_ = *δ*_*d*_ = 1, *K* = 10^5^, Φ_*G*_ = Φ_*S*_ = 0, and *U* = 0.1.

**Figure 5.**
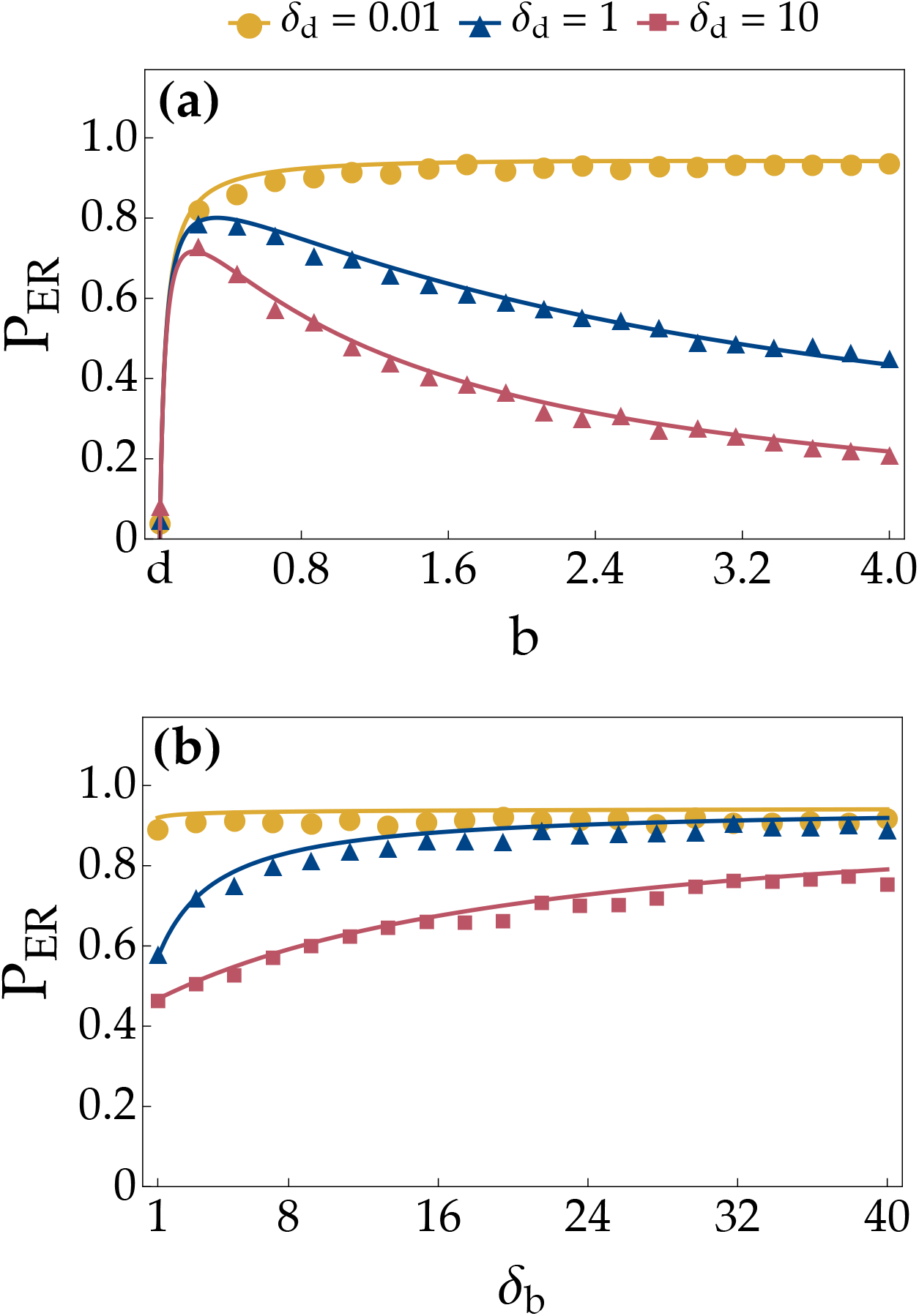
The probability of evolutionary rescue, *P*_*ER*_, as a function of (a) the resident density-independent birth rate, *b*, and (b) the sensitivity of residents to birth-limiting interactions, *δ*_*b*_, given different sensitivities of residents to death-promoting interactions, *δ*_*d*_. Lower resident birth rates appreciably enhance *P*_*ER*_ only if *δ*_*d*_ is sufficiently large. Otherwise, variance in mutant fitness is unaffected by changes in wild type density. Markers: proportion of 2 × 10^3^ replicate simulations that ended in mutant survival; solid curves: analytical approximation of *P*_*ER*_ using Eqs. (10) and (11). Unless stated otherwise, parameter values are *b* = 1, *d* = 0.03, *δ*_*b*_ = 3, *δ*_*d*_ = 1, *K* = 10^5^, Φ_*G*_ = 9, Φ_*S*_ = 0, and *U* = 0.1.

## 2 Model

In this section, an eco-evolutionary model of resident-invader dynamics is formulated and then modified for ER. The model’s constituent parameters and variables are listed in Table 1.

**Table 1:**
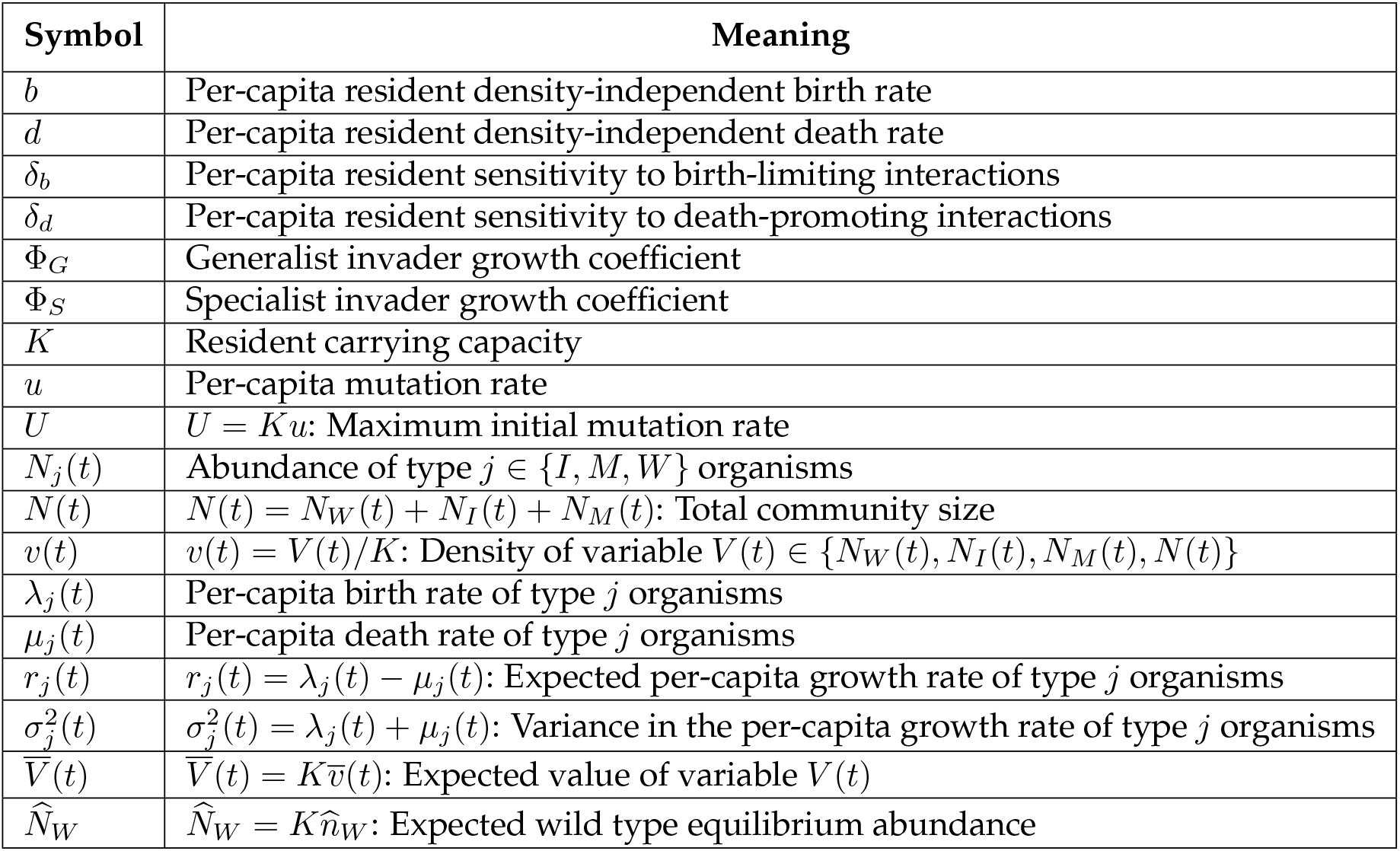
Model parameters and variables.

| Symbol | Meaning |
| --- | --- |
| $b$ | Per-capita resident density-independent birth rate |
| $d$ | Per-capita resident density-independent death rate |
| $\delta_b$ | Per-capita resident sensitivity to birth-limiting interactions |
| $\delta_d$ | Per-capita resident sensitivity to death-promoting interactions |
| $\Phi_G$ | Generalist invader growth coefficient |
| $\Phi_S$ | Specialist invader growth coefficient |
| $K$ | Resident carrying capacity |
| $u$ | Per-capita mutation rate |
| $U$ | $U = Ku$: Maximum initial mutation rate |
| $N_j(t)$ | Abundance of type $j \in \{I, M, W\}$ organisms |
| $N(t)$ | $N(t) = N_W(t) + N_I(t) + N_M(t)$: Total community size |
| $v(t)$ | $v(t) = V(t)/K$: Density of variable $V(t) \in \{N_W(t), N_I(t), N_M(t), N(t)\}$ |
| $\lambda_j(t)$ | Per-capita birth rate of type $j$ organisms |
| $\mu_j(t)$ | Per-capita death rate of type $j$ organisms |
| $r_j(t)$ | $r_j(t) = \lambda_j(t) - \mu_j(t)$: Expected per-capita growth rate of type $j$ organisms |
| $\sigma_j^2(t)$ | $\sigma_j^2(t) = \lambda_j(t) + \mu_j(t)$: Variance in the per-capita growth rate of type $j$ organisms |
| $\bar{V}(t)$ | $\bar{V}(t) = K\bar{v}(t)$: Expected value of variable $V(t)$ |
| $\hat{N}_W$ | $\hat{N}_W = K\hat{n}_W$: Expected wild type equilibrium abundance |

### 2.1 Eco-evolutionary model

Considered herein is a well-mixed community that changes according to a continuous-time birth–death process. Within the community are two species: a resident and an invader, together comprising *N* (*t*) ≥ 0 organisms. The resident species includes wild types and their mutant descendants—collectively referred to as residents—while the invader species contains organisms of a single type, referred to as invaders. The abundances of these three types are denoted by *N*_*W*_ (*t*) ≥ 0, *N*_*M*_ (*t*) ≥ 0, and *N*_*I*_ (*t*) ≥ 0, respectively, and so *N* (*t*) = *N*_*W*_ (*t*) + *N*_*M*_ (*t*) + *N*_*I*_ (*t*). Wild types are sensitive to interactions with invaders, which can be either generalists that proliferate independently of wild types or specialists that require wild types to proliferate. Mutants, in contrast, are resistant to interactions with invaders.

#### 2.1.1 Wild type dynamics

Residents have a soft carrying capacity of *K >* 0 such that, on average, *N*_*W*_ (*t*) ≤ *K*. The density of each variable *V* (*t*) ∈ {*N*_*W*_ (*t*), *N*_*I*_ (*t*), *N*_*M*_ (*t*), *N* (*t*)} with respect to *K* is denoted by *v*(*t*) = *V* (*t*)*/K*. To limit the number of model parameters, wild type fitness is assumed equally sensitive to intra- and interspecific interactions. Wild types give birth at a per-capita rate of *λ*_*W*_ (*t*) = max (0, *b* − *δ*_*b*_*n*(*t*)), where *b >* 0 is the per-capita resident density-independent birth rate and *δ*_*b*_ ≥ 0 is the per-capita sensitivity of residents to birth-limiting interactions. Similarly, wild types die at a per-capita rate of *µ*_*W*_ (*t*) = *d* + *δ*_*d*_*n*(*t*), where 0 *< d < b* is the per-capita resident density-independent death rate and *δ*_*d*_ ≥ 0 is the per-capita sensitivity of residents to death-promoting interactions.

#### 2.1.2 Invader dynamics

For simplicity, the rate of invader migration into the community is assumed low enough to have a negligible effect on invader dynamics. Furthermore, following previous theory, a worst-case scenario for wild types is considered, wherein invaders have a large enough carrying capacity to be unharmed by competition, and possess a negligible density-independent death rate (Longcamp & Draghi, 2025). These assumptions result in a per-capita invader death rate that is effectively zero (i.e., *µ*_*I*_ (*t*) = 0), ensuring that invaders avoid extinction and therefore drive wild types extinct. The per-capita invader birth rate is modeled as *λ*_*I*_ (*t*) = Φ_*G*_ + *n*_*W*_ (*t*)Φ_*S*_ ≥ 0, where Φ_*G*_ ≥ 0 is the per-capita wild type-independent invader growth rate— hereafter, the generalist invader growth coefficient—and Φ_*S*_ ≥ 0 is the specialist invader growth coefficient, defined as the average maximum value of the per-capita wild type-dependent invader growth rate, *n*_*W*_ (*t*)Φ_*S*_. Henceforth, focus is restricted to two alternative invader types: generalists with Φ_*G*_ *>* 0 and Φ_*S*_ = 0 and specialists with Φ_*G*_ = 0 and Φ_*S*_ *>* 0.

#### 2.1.3 Mutant dynamics

Wild types mutate into invader-resistant mutants at a per-capita rate of *u >* 0. Back mutations are ignored, and forward mutation events are assumed infrequent enough to have a negligible effect on the wild type growth rate (i.e., mutation events do not decrease *N*_*W*_ (*t*)). In the absence of invaders, mutants are assumed strongly deleterious and consequently rare. When invaders are present, however, mutant fitness is equal to that of wild types except mutants are unaffected by invader density and therefore have per-capita birth and death rates of *λ*_*M*_ (*t*) = max (0, *b* − *δ*_*b*_ (*n*_*W*_ (*t*) + *n*_*M*_ (*t*))) and *µ*_*M*_ (*t*) = *d* + *δ*_*d*_ (*n*_*W*_ (*t*) + *n*_*M*_ (*t*)), respectively, where hereafter, on average, *λ*_*M*_ (*t*) *>* 0.

#### 2.1.4 Expected community dynamics

In the present model, time advances in infinitesimal intervals, (*t, t* + *dt*). Consequently, at most one event occurs at a given moment, with the probability of simultaneous events being of negligible order, *O*(*dt*^2^). When an event *E* ∈ {*j* birth, *j* death, mutation} occurs, it changes a variable by an amount Δ_*E*_ ∈ {+1, −1}. Letting *P* (*E, N*_*j*_(*t*) + Δ_*E*_) denote the probability of type *j* ∈ {*W, I, M*} abundance, *N*_*j*_(*t*), undergoing state change Δ_*E*_ due to event *E*, the possible event probabilities are

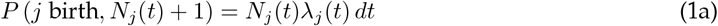

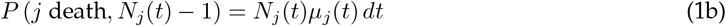

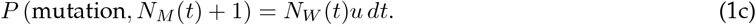

Since the birth and death probabilities in Eq. (1) scale with *N*_*j*_(*t*), when *N*_*j*_(*t*) is large, the total demographic change in *N*_*j*_(*t*) over a given finite time interval can be viewed as the sum of many small state changes, each of magnitude 1 ≪ *N*_*j*_(*t*); and by the same reasoning, assuming hereafter that *N*_*W*_ (*t*) ≫ 1, the change in *N*_*M*_ (*t*) caused by mutations can likewise be treated as the accumulation of many small contributions. Accordingly, the infinitesimal change in a large *N*_*j*_(*t*) value, denoted *dN*_*j*_(*t*), can be approximated as a normally distributed random variable, 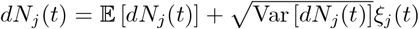, where 𝔼 [*dN*_*j*_(*t*)] and Var [*dN*_*j*_(*t*)] are the mean and variance of *dN*_*j*_(*t*), respectively, and *ξ*_*j*_(*t*) is a standard normal variable. When applied to each *N*_*j*_(*t*), this widely used *diffusion approximation* yields

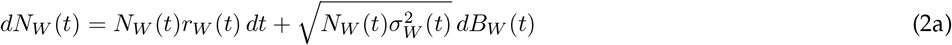

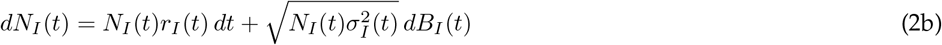

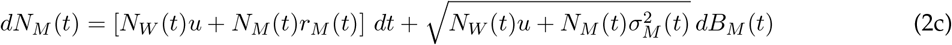

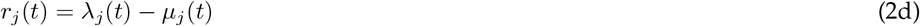

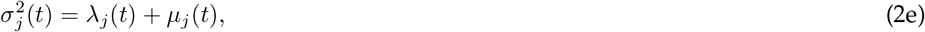

where *B*_*W*_ (*t*), *B*_*I*_ (*t*), and *B*_*M*_ (*t*) are independent standard Brownian motions, changing in respective in-crements of 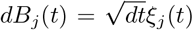; the variable *r*_*j*_(*t*) represents type *j*’s expected per-capita growth rate; and *σ*^2^(*t*) denotes the variance in type *j*’s per-capita growth rate (Czuppon & Traulsen, 2021; Ethier & Kurtz, 1986, Chapter 11). The second term of each differential equation in Eq. (2) quantifies the change in *N*_*j*_(*t*) caused by stochastic fluctuations in birth, death, and mutation rates, while the first term gives the deterministic change that would occur without such stochasticity. Since the ratio of the stochastic term to the deterministic term scales as 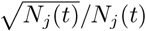, it approaches zero as *N*_*j*_(*t*) increases. Therefore, once *N*_*j*_(*t*) is sufficiently large, variance in *dN*_*j*_(*t*) becomes negligible relative to the mean, and *dN*_*j*_(*t*) is consequently well-approximated by its expectation:

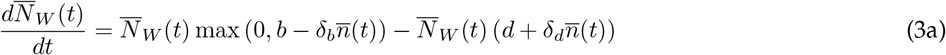

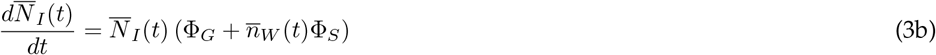

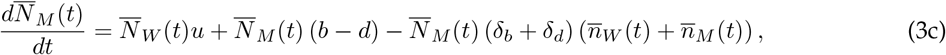

with 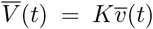 denoting the expected value of variable *V* (*t*). Letting 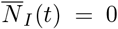 and 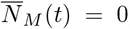, and then solving 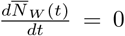 for 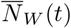, reveals that Eq. (3) exhibits an equilibrium at which wild type abundance is

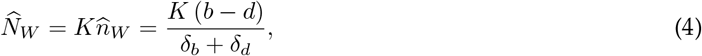

where henceforth the total per-capita sensitivity of residents to biological interactions is *δ*_*b*_ + *δ*_*d*_ ≥ *b* − *d*, so that 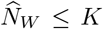. Since the derivative of 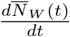 with respect to 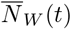 has a negative value of − (*b* − *d*) when evaluated at 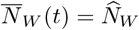, the equilibrium is locally stable throughout parameter space. Thus, when invaders and mutants are vanishingly rare, the wild type population is expected to return toward 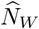 following a small fluctuation in size.

### 2.2 Evolutionary rescue

Figure 1 illustrates the dynamics of the eco-evolutionary model, modified for ER from invasion. At times *t <* 0, invaders are absent from the community. In contrast, *K* is assumed large such that wild types have an abundance of 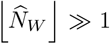, where the floor function, 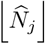, ensures a count of whole organisms. Meanwhile, following previous theory, a single mutant is assumed present as standing variation (e.g., Czuppon et al., 2023; Marrec & Bank, 2023). At time *t* = 0, an invader appears, giving the community an initial composition of 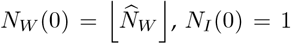, and 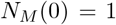. The wild type population then declines as invaders proliferate. This decline is faster when invaders are generalists (Figure 1a) instead of specialists (Figure 1b), because the specialist growth rate declines with wild type density. Since ER is nearly certain under sufficiently high mutation rates (e.g., Marrec & Bitbol, 2020), *u* is set to *u* = *U/K* so that the mutation rate, *N*_*W*_ (*t*)*u* = *n*_*W*_ (*t*)*U*, has a maximum initial value of *U* = *Ku <* 1 and is thus, on average, less than one. Mutants are considered established, and ER is said to occur, only if 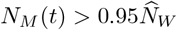.

## 3 Methods

An analytical approximation for the probability of ER, *P*_*ER*_, is derived and its predictions are tested with stochastic simulations. The first part of this section presents the building blocks of the approximation; the second outlines the simulation algorithm.

### 3.1 Analytical building blocks

For analytical simplicity, ER is classified as arising from *de novo* mutation only if the mutant lineage from standing variation is extinct when the ER process completes; otherwise a successful ER event is attributed to standing variation. Under this assumption, the two possible routes to ER—standing variation and *de novo* mutation—are mutually exclusive. Therefore, by the law of total probability:

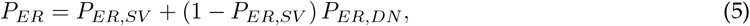

where *P*_*ER,SV*_ is the probability of ER via standing variation and *P*_*ER,DN*_ is the probability of ER via *de novo* mutation. To approximate *P*_*ER,SV*_ and *P*_*ER,DN*_, resident carrying capacity is assumed large (i.e., *K* ≫ 1) such that, when wild types are present at a non-negligible density, their abundance changes deterministically (i.e., according to Eq. (3)) and thus 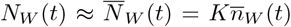. Invader abundance is similarly approximated as 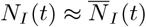. Furthermore, for the window of time relevant to mutants escaping stochastic extinction, mutant density, *n*_*M*_ (*t*), is assumed low enough for the per-capita mutant birth and death rates to be 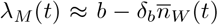 and 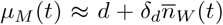, respectively. Under the latter assumption, mutants reproduce and die independently of one another, meaning that *P*_*ER,SV*_ can be approximated using time-inhomogeneous branching process theory (Kendall, 1948; Uecker & Hermisson, 2011). By plugging *λ*_*M*_ (*t*) and *µ*_*M*_ (*t*) into the extinction probability derived by Kendall (1948, Eq. (18) therein), one obtains

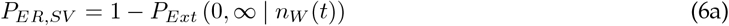

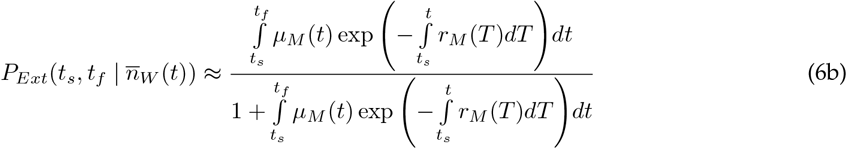

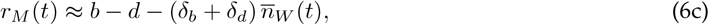

where 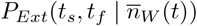 is the probability that a mutation present as a single copy at some starting time, *t*_*s*_ ≥ 0, is extinct by some final time, *t*_*f*_ *> t*_*s*_, conditional on 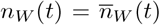 for all *t* ∈ [*t*_*s*_, *t*_*f*_]. Assuming mutations are also rare enough to appear independently of one another at any given time *t* = *t*_*M*_ *>* 0, the number of times a *de novo* mutant lineage appears and establishes is a Poisson-distributed random variable. The expectation of this random variable is 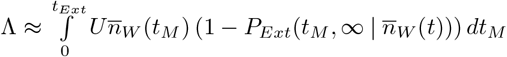, where 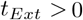 is the time of wild type extinction. Thus, by taking the complement of the Poisson extinction probability, Λ^0^ exp (−Λ) */* (0!), one attains

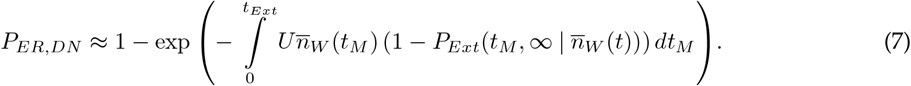

Both Eqs. (6) and (7) require an approximation of 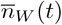 and hence 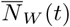. Readers uninterested in the derivation of 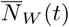 can safely skip to *Simulations*.

#### 3.1.1 Approximation of 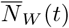

The approach adopted herein to approximate 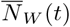 differs depending on whether the per-capita invader growth rate is large, *r*_*I*_ (*t*) ≫ 0, or small, *r*_*I*_ (*t*) ≈ 0, and therefore whether invasion is fast or slow. In both cases, the per-capita wild type birth rate is assumed nonzero (i.e., 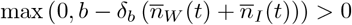) while 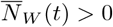.

##### Approximation of 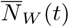 under fast invasion

For the case of *r*_*I*_ (*t*) ≫ 0, invasion is assumed fast enough such that 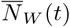 is non-negligible only during the brief moment when 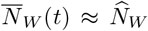 . The dynamics of 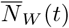 during that brief moment are captured by first supposing 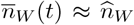 so that 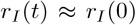, and therefore 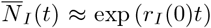. After substituting 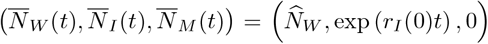 into the right-hand side of Eq. (3a), one can then solve Eq. (3a), obtaining 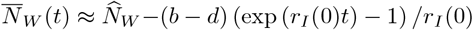. Assuming *r*_*I*_ (0) is large enough for exp (*r*_*I*_ (0)*t*) − 1 ≈ exp (*r*_*I*_ (0)*t*), this becomes

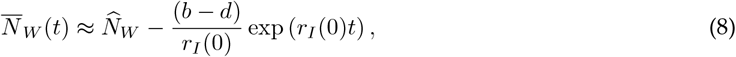

which gives a wild type extinction time of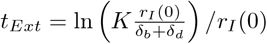

##### Approximation of 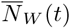 under slow invasion

For the case of *r*_*I*_ (*t*) ≈ 0, first considered are the dynamics of the total community, 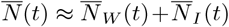, while it is near its initial value of 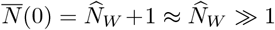. Letting 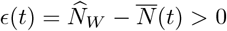 represent a small reduction in 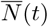 from 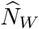, one finds 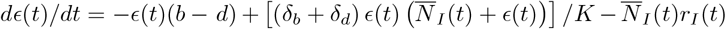, which represents the rate at which the distance between 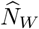 and 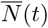 shrinks (indicated by *dϵ*(*t*)*/dt <* 0) or expands (indicated by *dϵ*(*t*)*/dt >* 0) following a small perturbation. For simplicity, one can linearize *dϵ*(*t*)*/dt* around *ϵ*(*t*) = 0, acquiring 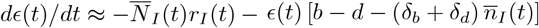. With *r*_*I*_ (*t*) ≈ 0, this rate becomes positive only once invaders surpass a critical abundance of 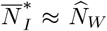, indicating that (i) 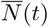 stays close to 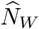while 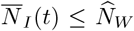, and thus (ii) wild types have an abundance of 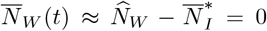 once 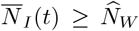. Accordingly, for times when 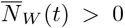, the community is assumed to maintain an effectively constant size of 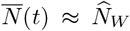, which allows one to approximate 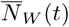 by calculating 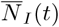 and then rearranging 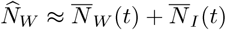.

For generalist invaders with a per-capita growth rate of 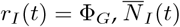 is acquired by solving Eq. (3b) with 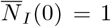, yielding 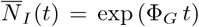. In contrast, for specialist invaders with 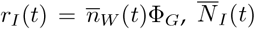 is attained through the following steps. First, let 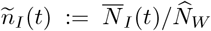represent invader density relative to the assumed maximum community size, 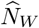. Second, simplify Eq. (3b) with 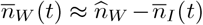 and 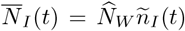, obtaining 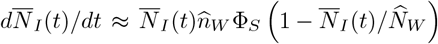 . Third, solve the latter rate with 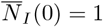, which gives 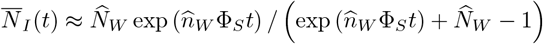 . Finally, simplify 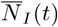 by supposing 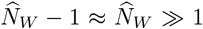, such that 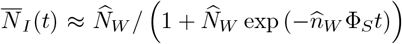 . Rearranging 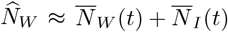 then gives

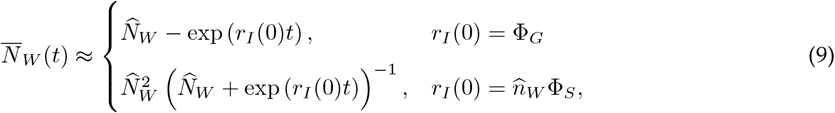

which yields a wild type extinction time of 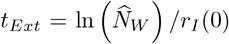 when *r*_*I*_ (0) = Φ_*G*_ and *t*_*Ext*_ → ∞ when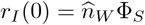.

### 3.2 Simulation algorithm

Analytical predictions are tested with simulations of a tau-leaping algorithm (Cao et al., 2006). A simulation is initialized with 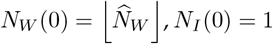, and *N*_*M*_ (0) = 1. Each variable, *N*_*j*_(*t*), then changes according to its corresponding events (*E*), state changes (Δ_*E*_), and transition rates (*P* (*E, N*_*j*_(*t*) + Δ_*E*_) */dt*) listed in Eq. (1). The size of each tau-leap, and the number of times each event occurs during the leap, are computed according to the procedure outlined in Sections IIC (excluding Step 3) and IV of Cao et al. (2006). As implemented, this procedure ensures that, if *N*_*j*_(*t*) *< N*_*c*_ = 10, then all transition rates corresponding to a nonzero state change in *N*_*j*_(*t*) are updated according to a nearly exact version of the Gillespie algorithm (Gillespie, 1976; see parameter *n*_*c*_ = 10 in Cao et al., 2006). Moreover, the algorithm helps prevent *N*_*j*_(*t*) from becoming negative during a leap by approximately bounding the per-leap change in each of *N*_*j*_(*t*)’s transition rates, relative to the sum of all transition rates, by a value of *ρ* = 0.03 (see parameter *ϵ* = 0.03 in Cao et al., 2006). If *N*_*j*_(*t*) becomes negative despite this precaution, then it is set to *N*_*j*_(*t*) = 0. An undefined leap size is avoided by adding 10^−9^ to the output of Cao et al.’s (2006) Eq. (32a) if the output equals zero, and the same modification is made to their Eq. (32b) if its output equals zero. A simulation proceeds until either *N*_*W*_ (*t*) = *N*_*M*_ (*t*) = 0 or 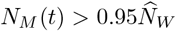 . All numerical calculations and simulations are performed using Mathematica 13.2.0.0.

## 4 Results

In this section, the probability of ER via standing variation, *P*_*ER,SV*_, and that via *de novo* mutation, *P*_*ER,DN*_, are analytically approximated. Then, the total ER probability, *P*_*ER*_ = *P*_*ER,SV*_ +(1 − *P*_*ER,SV*_) *P*_*ER,DN*_, is analyzed. For concision, the terms “expected” and “per-capita” are omitted in references to quantities within Table 1. The derivations of *P*_*ER,SV*_ and *P*_*ER,DN*_ are presented with a focus on biological assumptions, with the corresponding mathematical details relegated to Appendices A and B, respectively. Any simplifying assumptions introduced hereafter apply only to these derivations, which are made using Eqs. (8) and (9) for wild type abundance, 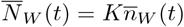; Eqs. (6) and (7) for *P*_*ER,SV*_ and *P*_*ER,DN*_; and thus Eq. (6b) for the probability that a mutation present at time *t*_*s*_ *>* 0 is extinct by some time 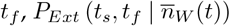.

### 4.1 The probability of evolutionary rescue via standing variation

As shown in Figure 1, 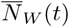 remains near its initial value, 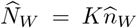, until invaders become sufficiently abundant. Meanwhile, the first mutant in the community—the standing variant—is nearly neutral, with a growth rate of 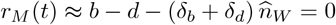. For this reason, *P*_*ER,SV*_ is approximated under the assumption that wild type density is 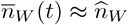, and hence the standing variant is neutral, until some critical time, *t*_*c*_ *>* 0. Since a neutral lineage is certain to go extinct given enough time (i.e., 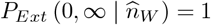; Kendall, 1948), attention is restricted to regimes where invaders quickly displace wild types (i.e., Eq. (8) is used for 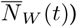, causing the standing variant to be only briefly neutral. In particular, the initial invader growth rate, *r*_*I*_ (0), is assumed large enough that not only is *t*_*c*_ effectively equal to the wild type half-life, 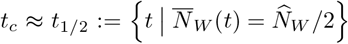, but also wild types become vanishingly rare, 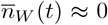, once *t > t*_1*/*2_. Under these assumptions, Eq. (6a) becomes 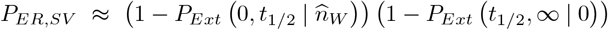, which simplifies to

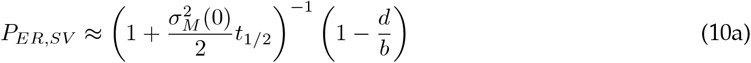

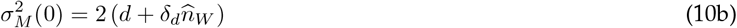

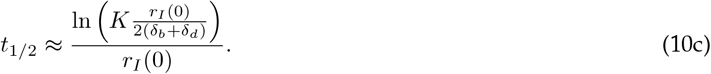

Equation (10) says that, at least approximately speaking (assumed hereafter)—and all else equal (also assumed henceforth)—*P*_*ER,SV*_ can be increased by reducing any of three key quantities. The first is the initial variance in mutant fitness, 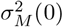, specifically the half attributable to random deaths: reducing this variance diminishes early stochastic declines in mutant abundance, helping the standing variant persist while 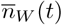 is high and competition between mutants and wild types is consequently strong. The second is *t*_1*/*2_: a shorter *t*_1*/*2_ means wild types go extinct sooner, relieving the standing variant of competition more quickly. The third is the density-independent probability of mutant extinction, *d/b*: even after wild types are extinct, mutants must have a sufficiently positive density-independent growth rate, *r*_*M*_ = *b*−*d*, to avoid stochastic extinction. If a parameter shift does not directly reduce one of these three quantities, the shift may still indirectly increase *P* by lowering 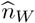 or raising *r*_*I*_ (0). A higher *r*_*I*_ (0) accelerates the wild type decline and hence reduces *t*, while a lower 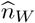 primarily decreases 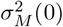. A decrease in 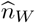 can also diminish *t*_1*/*2_, specifically through a reduction in the total sensitivity of residents to biological interactions, *δ*_*b*_+*δ*_*d*_, because that sensitivity—along with *r*_*I*_ (0)—governs how rapidly wild types are displaced. However, given the large resident carrying capacities, *K* ≫ 1, considered herein, the logarithmic term in *t*_1*/*2_ varies only weakly with *δ*_*b*_ + *δ*_*d*_, rendering 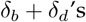 effect on *t*_1*/*2_ negligible compared with its effect on 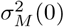.

### 4.2 The probability of evolutionary rescue via *de novo* mutation

In contrast to *P*_*ER,SV*_, *P*_*ER,DN*_ is approximated with a focus on regimes where the wild type population declines slowly, and *r*_*I*_ (0) is therefore small, giving *de novo* mutations time to appear (i.e., Eq. (9) is used for 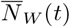. The magnitude of *r*_*I*_ (0) is assumed so small that, throughout the establishment of a focal mutant lineage appearing at time *t*_*M*_, wild types maintain an effectively constant density of 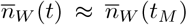. Using this constant density, Eq. (7) is first solved, and then the solution is simplified by taking its limit as *K* → ∞, yielding

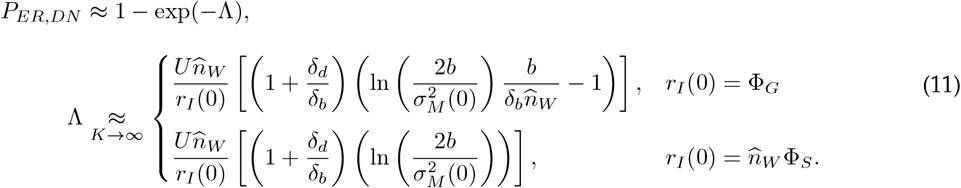

In Eq. (11), the factor 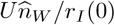 comes from integrating the mutation rate, 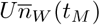, across all possible mutation times *t*_*M*_, while the bracketed term accounts for the probability that a mutant lineage establishes. Since the right-hand side of Eq. (11) increases with both terms, the equation shows that *P*_*ER,DN*_ rises with the cumulative density of wild types across all *t*_*M*_, as more wild types implies more ER opportunities. This means that *P*_*ER,DN*_ not only grows with a smaller *r*_*I*_ (0)—due to the slower wild type decline—but is also larger when invaders are specialists with 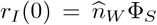, rather than generalists with *r*_I_ (0) = Φ_*G*_. The bracketed term is larger with specialists because the specialist growth rate, 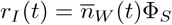, declines with 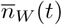. Similarly, for a given invader growth coefficient, Φ = Φ = Φ, the factor 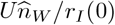 is larger with specialists because 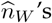 effect on the initial mutation rate, 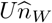, is offset by its corresponding effect on 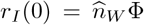. Thus, with specialists, a lower 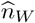 can increase *P*_*ER,DN*_ by boosting the total chance of mutant establishment, whereas with generalists a reduction in 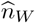 always lowers *P*_*ER,DN*_ through adecrease in mutational input.

### 4.3 The probability of evolutionary rescue

The behavior of *P*_*ER*_ = *P*_*ER,SV*_ + (1 − *P*_*ER,SV*_) *P*_*ER,DN*_ is now analyzed. All results are interpreted in light of the analytical approximation, derived using Eqs. (10) and (11).

#### 4.3.1 Rescue is most likely with slowly invading specialists or rapidly invading generalists

The *P*_*ER*_ approximation reveals that the likely route to ER—standing variation or *de novo* mutation—differs based on the initial invader growth rate, 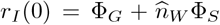, where Φ_*S*_ and Φ_*G*_ are the generalist and specialist invader growth coefficients, respectively, and 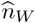 is the initial wild type density, 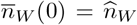. If *r*_*I*_ (0) is sufficiently small, ER is more likely to occur through *de novo* mutation than standing variation (Figure 2a). Slower invasion leads to a slower decline in wild type density, 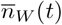, creating a longer time interval wherein a new mutation can arise. If *r*_*I*_ (0) is adequately large, on the other hand, ER is most likely to occur through standing variation (Figure 2a). By reducing the wild type half-life, a higher *r*_*I*_ (0) lowers the effect of 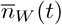 on the mutant birth rate, 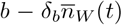, and death rate, 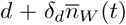, which allows the lineage from standing variation to achieve a non-negligible competitive advantage over wild types at an earlier time and thus diminishes the lineage’s extinction probability.

Broadly, this effect of *r*_*I*_ (0) on the likely route to ER means that *P*_*ER*_ is highest when invasion is either sufficiently slow or sufficiently fast (Figure 2a). In terms of invader type, it also means that, for a given Φ_*S*_ = Φ_*G*_ = Φ, *P*_*ER*_ is highest when invaders are either slowly invading specialists with 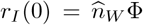 or rapidly invading generalists with *r*_*I*_ (0) = Φ (Figure 2b). In the former case, specialists are favorable because they sometimes fail to encounter their victims, providing additional time for *de novo* mutations to arise; in the latter, generalists are favorable because their constant growth rate allows standing variants to be relieved from competition more rapidly. The establishment of standing variation can also be facilitated by an increase in Φ_*S*_, but the resulting boost in *P*_*ER*_ is less pronounced than for generalists (Figure 2b).

#### 4.3.2 Lower resident density-independent birth rates can facilitate rescue from generalists

Interestingly, *P*_*ER*_ can vary non-monotonically with the resident density-independent birth rate, *b* (Figure 3a). As *b* rises from near the resident density-independent death rate, *d, P*_*ER*_ initially increases (Figure 3a), because mutants, whether from standing variation or *de novo* mutation, require a sufficiently large density-independent growth rate, *r*_*M*_ = *b* − *d*, to escape stochastic extinction. Once *b* is adequately larger than *d*, however, the qualitative behavior of *P*_*ER*_ depends on the initial invader growth rate, *r*_*I*_ (0). If *r*_*I*_ (0) is small enough to favor *de novo* mutation as the likely route to ER, *P*_*ER*_ continues increasing with *b*, irrespective of whether invaders are generalists with *r*_*I*_ (0) = Φ_*G*_ (Figure 3a) or specialists with 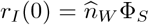 (Figure 3b). For the generalist case, this increase in *P*_*ER*_ occurs because a higher *b* not only increases *r*_*M*_ but also augments the initial wild type density, 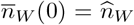, which raises the initial mutation rate, 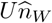 . For the specialist case, the boost in 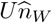 is offset by a decline in 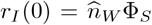, so *P*_*ER*_ increases with *b* due solely to the higher *r*_*M*_ .

By contrast, if *r*_*I*_ (0) is large enough to favor standing variation, reducing *b* can increase *P*_*ER*_ (Figure 3a). In such a case, the corresponding decrease in 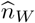 lowers the initial variance in mutant fitness, 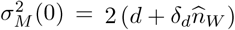, ensuring that mutants experience fewer random deaths and are consequently less likely to go extinct. With generalists, this benefit of reducing *b* can suffice to raise *P*_*ER*_ (Figure 3a); but with specialists, *P*_*ER*_ tends to diminish with *b* (Figure 3b), because the smaller 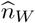slows wild type displacement enough for the cost of a longer wild type half-life—and thus prolonged competition between mutants and wild types— to outweigh the benefit of lower mutant variance. Although increasing *d* also lowers 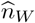, it differs from reducing *b* in that it directly raises mutant variance, leading to more random mutant deaths. Consequently, *P*_*ER*_ consistently decreases with increasing *d*, regardless of whether invaders are generalists (Figure 3c) or specialists (Figure 3d).

#### 4.3.3 Birth-limiting competition can aid rescue from both generalists and specialists

Another factor with which *P*_*ER*_ can vary non-monotonically is the sensitivity of residents to birth-limiting interactions, *δ*_*b*_ (Figure 4a). With generalist invaders, the effect of *δ*_*b*_ on *P*_*ER*_ resembles that of the resident density-independent birth rate, *b*: if generalists invade slowly, a larger *δ*_*b*_ hinders ER (Figure 4a) by lessening the initial wild type density, 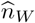, and hence mutational input, whereas if they invade quickly, a greater *δ*_*b*_ facilitates ER (Figure 4b) by shrinking the initial mutant variance, 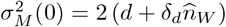, through . In contrast to a lower *b*, however, a higher *δ_b_* can notably enhance *P*_*ER*_ when invaders are specialists (Figure 4b). This occurs because, unlike the fitness effect of density-independent factors, the reduction in mutant fitness caused by birth-limiting competition decays with 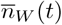, which allows the benefit of lower mutant variance to more readily outweigh the cost of lower mutant fitness. The reduced variance can then enhance *P*_*ER*_, since 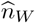has a neutral effect on mutational input when invaders are specialists. This enhancement is most pronounced when specialist invasion is slow—but not so slow that ER is effectively certain (Figure 4b)—because then mutations tend to appear *de novo* and can thus appear at relatively late times when competition is more relaxed. If invasion is instead too fast, then, as with a lower *b*, a higher *δ*_*b*_ can hinder ER (Figure 4b) by prolonging competition between wild types and standing variants.

Although the effect of death-promoting competition on mutant fitness also decays with 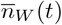, a stronger resident sensitivity to death-promoting interactions, *δ*_*d*_, does not facilitate ER, neither from generalists (Figure 4c) nor from specialists (Figure 4d). Instead, *P*_*ER*_ increases with a reduction in *δ*_*d*_, because as *δ*_*d*_ declines, the influence of 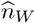 on mutant variance weakens, causing such variance to become increasingly insensitive to changes in wild type density. Importantly, this means that, when *δ*_*d*_ is sufficiently small, *P*_*ER*_ can no longer appreciably increase with a reduction in *b* (Figure 5a) or a boost in *δ*_*b*_ (Figure 5b), since the accompanying decrease in wild type density no longer meaningfully lowers mutant variance.

## 5 Discussion

Herein, an analytical approximation has been derived for the probability of ER, *P*_*ER*_ (Eqs. (10) and (11)), from an antagonist invasion. The approximation reveals that *P*_*ER*_ peaks when invaders are either slowly invading specialists with a growth rate that diminishes as wild types decline or rapidly proliferating generalists with a constant growth rate, as the former allows more time for *de novo* mutations to arise while the latter quickly releases standing variants from competition with wild types (Figure 2). The main result is that a reduction in the resident birth rate—but not an increase in the death rate—can enhance *P*_*ER*_ (Figure 3) by lowering wild type density and, in turn, weakening the effect of death-promoting competition on the variance in mutant fitness. Both a lower birth rate and a higher death rate decrease wild type density, but only the former lessens the total rate of random mutant deaths and therefore mutant variance. If the lower birth rate is density-independent, it notably increases *P*_*ER*_ only when invaders are generalists (Figure 3), because, with specialists, fewer resident births slow invasion and thus strengthen the competitive effect of wild types on mutants. In contrast, if the lower birth rate stems from birth-limiting competition, the decreased variance can appreciably raise *P*_*ER*_ whether invaders are generalists or specialists (Figure 4), since the intensity of such competition declines with wild type density, allowing mutants to maintain a sufficiently large growth rate regardless of invader type. Even when competition-induced, however, a lower resident birth rate promotes ER only if mutants are sufficiently sensitive to death-promoting competition (Figure 5), ensuring that mutant variance depends on wild type density.

### Antagonistic interactions and victim specificity

Among existing ER models with antagonistic interactions (e.g., Shang et al., 2024; Vanselow et al., 2022; van Velzen, 2023; Yamamichi & Miner, 2015), those that consider differences in victim specificity typically focus on ER from abiotic stress, with adaptation occurring through quantitative trait evolution (Osmond et al., 2017; van Velzen, 2023). This contrasts with the present model, wherein ER proceeds via a single mutation conferring resistance to biological invasion. Nonetheless, the present work qualitatively extends at least two such quantitative genetic studies by identifying additional ways in which victim specificity can shape ER dynamics. In particular, simulations by van Velzen (2023) showed that specialist predators— but not generalists—induced a form of indirect ER (Hermann & Becks, 2022; Yamamichi & Miner, 2015), whereby predator evolution toward greater specificity for less-defended prey reduced predation on more-defended prey enough to prevent extinction. The present study complements this result by demonstrating that victim specificity can also promote direct ER driven by adaptation to antagonists themselves: when specialized antagonists have sufficiently low growth coefficients, their fitness declines sharply as suitable victims become rare, buying time for resistance mutations to arise.

Using a quantitative genetic model, Osmond et al. (2017) similarly showed that greater prey specificity among predators can promote direct prey ER by causing predator fitness to decline with prey density. In their model, this fitness decay limits prey mortality under intense predation, allowing prey populations to better track an abiotically shifting phenotypic optimum (Osmond et al., 2017). When predation intensity is low, however, generalist predators are favorable, as their time-invariant densities ensure stronger effects on prey mortality that produce a larger reduction in birth-limiting competition among prey, thereby increasing prey fecundity and, consequently, accelerating prey adaptation (Osmond et al., 2017). The present results are consistent with these findings in that antagonist victim specificity can either promote or hinder ER depending on ecological conditions, which here are determined by the antagonist growth coefficient. If that coefficient is sufficiently high so that even specialists rapidly displace their victims, then generalists are favorable, because their steadier proliferation induces a quicker victim displacement that, in turn, rapidly relieves antagonist-resistant standing variants of competition with their wild type ancestors. This benefit of reduced competition, known as competitive release, has been observed in both laboratory experiments (Pena-Miller et al., 2013) and theoretical work (Uecker et al., 2014) without invasion-induced wild type decline.

### Specialist invasion and the wild type decline

Similar to the present work, other theoretical studies of ER have considered invasion as the primary cause of wild type decline (DeLong & Belmaker, 2019; Golas et al., 2021; Jiao et al., 2020; Longcamp & Draghi, 2025; Van Dyken, 2020). Although most of these models do not account for differences in invader victim specificity, they suggest—as highlighted herein—that specialist invasion can alter ER dynamics by inducing a dynamic coupling between the rate of wild type decline and the process driving that decline. In simulation experiments by DeLong and Belmaker (2019), for instance, the probability of ER from invasive predators increased when predation drove prey evolution toward a smaller body size that, in addition to boosting prey fecundity and lessening the predator attack rate, reduced the energetic value of prey, thereby slowing invasion. Moreover, in simulations of ER from pathogen invasion, Jiao et al. (2020) observed that pathogen-induced mortality could reach levels that so strongly depressed wild type density that disease prevalence fell below a critical reproductive threshold, ending the outbreak and obviating the need for ER.

Another example of how a wild type–invader coupling can shape ER dynamics was highlighted by Longcamp and Draghi (2025), who analyzed ER from specialist invaders exploiting resident-constructed habitats. Their model is nearly identical to the specialist invader model herein, except they ignored birth-limiting interactions and assumed invader proliferation depends on habitat rather than wild type density (Longcamp & Draghi, 2025). Under these assumptions, they derived an approximation for ER via *de novo* mutation, 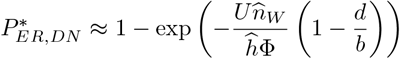, and showed that residents can reduce extinction risk by lowering initial habitat density, 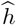, thereby slowing invasion enough to buy time for mutation. The present study therefore extends this result by showing that a similar time-buying effect can arise when residents themselves are the exploited resource. To enable direct analytical comparison, one can take the limit of Eq. (11) as birth-limiting competition vanishes (i.e., *δ*_*b*_ → 0), which yields the ER probability analogous to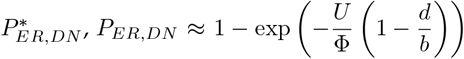. The key difference between 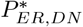 and *P*_*E R,DN*_ is that, unlike the mutational input term in 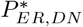(i.e., 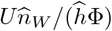), that in *P*_*ER,DN*_ (i.e., *U/*Φ) is independent of initial wild type density, 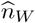, because the time-buying benefit of slower specialist invasion exactly offsets the mutational cost of fewer wild types. This distinction has two implications: ER from direct exploitation can be much less likely than from habitat exploitation when habitats are sufficiently ephemeral to impede specialists, yet the chance of ER from specialists can exceed that from generalists with constant growth rates, because only for specialists does the wild type–invader coupling counterbalance the cost of reduced mutational input.

Other studies have similarly shown that a wild type–invader coupling can neutralize the effects of key quantities on ER (Golas et al., 2021; Van Dyken, 2020), eliminating costs that often emerge without invasion-driven decline (e.g., Martin et al., 2013; Orive et al., 2019; Uecker et al., 2014). In particular, Golas et al. (2021) observed in simulations of pathogen-driven ER that a lower initial resident abundance—despite reducing the initial mutation rate—did not correlate with the probability of ER, due to a corresponding slowdown of invasion. Similarly, Van Dyken (2020) showed that when invaders are spatially spreading interference competitors, the probability of ER is independent of the invader dispersal rate, because the reduced mutational input associated with faster wild type decline is counterbalanced by a wider range of spatial locations where invader-resistant mutants have a selective advantage. Thus, the present work contributes to a growing taxonomy of predicted mechanisms by which the dependence of specialist invaders on wild type density can alter ER dynamics. An example of ER in which such dependence could have influenced mutational input is the ER of the Kauaian field cricket population, which adapted to a highly specialized parasitoid through what was possibly a *de novo* mutation (Tinghitella, 2008; Zhang et al., 2021; Zuk et al., 2006).

### Resident life history and mutant establishment

Across species, life histories often fall along a slow-fast continuum, with slow species characterized by low birth and death rates and fast species by high birth and death rates (Hernández-Yáñez et al., 2022). The present work connects with theory showing that low birth rates and high death rates—shared by both mutant and wild type residents—can differentially affect the probability of mutant establishment during ER (Czuppon et al., 2023; Raatz & Traulsen, 2023; Vinton & Vasseur, 2020). Czuppon et al. (2023) showed that ER is more likely under intraspecific birth-limiting competition than death-promoting competition, because the latter generates greater variance in mutant fitness. Similarly, Vinton and Vasseur (2020) observed that simulated populations persisted longer when environmental stress reduced birth rates rather than increased death rates. Extending these results, the present work reveals that a lower resident birth rate is not only less harmful than a higher resident death rate but can also facilitate ER relative to a higher birth rate by reducing the variance in mutant fitness generated by death-promoting competition. This prediction is supported by empirical studies indicating that herbaceous perennial species with low juvenile survival and bird species with low mean per-capita recruitment across life cycles are less vulnerable to extinction than higher-birth-rate counterparts (Hernández-Yáñez et al., 2022); and that birds often experience strong density-dependent mortality (Sæther et al., 2016), with demographic stochasticity being lowest in species with slow life histories (Sæther et al., 2004).

A closely related result was reported by Raatz and Traulsen (2023), who observed in simulations of a quantitative genetic model that the probability of ER increased with reductions in the resident birth rate. They suggested that this increase was likely driven by higher mutant survival probability, but could also have arisen because higher birth rates caused the ancestral lineage to decline more rapidly, thereby decreasing total mutational input (Raatz & Traulsen, 2023). The present study complements their work by showing analytically that, even when higher resident birth rates increase mutational input, lower birth rates can nonetheless promote ER by increasing mutant establishment probability. Moreover, whereas Raatz and Traulsen’s (2023) model—as well as non-ER theory highlighting a variance-diminishing benefit of lower birth rates (Parsons et al., 2018)—does not include death-promoting competition, the present work reveals that when such competition is present, lower birth rates can promote ER indirectly by weakening the effect of wild type density on mutant variance, even when they do not directly facilitate ER.

In the present invasion framework, birth-limiting competition is a more robust facilitator of ER than a lower density-independent birth rate, because it imposes a weaker cumulative reduction in mutant fitness that is more easily offset by reduced mutant variance. A lower resident density-independent birth rate only appreciably aids ER when invaders are generalists, mutations arise from standing variation, and mutant fitness remains sufficiently high. By contrast, lower variance from birth-limiting competition can notably promote ER from generalists via standing variation without constraint, and for specialists it can facilitate ER via both stand variation and *de novo* mutations, with the effect being stronger for *de novo* because they appear later when competition is weaker. Under all other conditions, however, lower resident birth rates tend to hinder ER by reducing mutational input and mutant fitness, consistent with previous theory (Marrec & Bank, 2023; Nyhoegen et al., 2024).

### Model shortcomings and future theory

Throughout this study, several assumptions have been made that could be worth relaxing in future work. For example, the model assumes that wild types are equally sensitive to intra- and interspecific interactions. As a result, for low-to-moderate invader growth rates where wild type extinction is not effectively instantaneous, invasion proceeds largely via replacement: each additional invader reduces wild type fitness about as much as an extra resident, keeping total community size nearly constant until wild types are extinct. An important consequence of this constant size is that wild type birth rates remain nonzero throughout most of their decline. In turn, the model does not recover a result of previous theory showing that, because birth rates have a minimum of zero while death rates can grow without bound (see also Draghi et al., 2024), the latter can generate notably higher ER probabilities by enabling faster competitive release for standing variation (Czuppon et al., 2023). Extending the model to include novel per-capita invader effects may support this prediction, revealing that ER from death-promoting invaders such as predators can be more likely than ER from birth-limiting invaders such as brood parasites. However, the opposite ranking may emerge for specialist invaders whose growth rate declines sharply with wild type fitness.

Two other notable assumptions of the present model are that, on average, an invader immediately harms wild types upon contact and simultaneously gains benefits from the interaction. Relaxing these assumptions could appreciably extend wild type extinction times—for example, through longer predator handling times (Abrams, 1990) or latent infection stages (Gandon, 2016)—thereby increasing the time available for resident mutation while also preventing mutant competitive release. An especially interesting extension could involve relaxing two additional assumptions: negligible invader mortality and complete mutant resistance. When these assumptions are relaxed, mutant and specialist invader abundances may oscillate, with mutants experiencing positive growth only when rare enough for specialists to die more frequently than they reproduce. Such dynamics could put mutants at risk of short-term post-ER extinction by driving them to low abundances shortly after establishment (Longcamp & Draghi, 2023). How factors such as invader handling time and stage structure differentially affect the total probability of ER and persistence through oscillations remains unclear.

Moreover, also assumed in the present model is that a single standing variant is present at the onset of invasion. To see when this condition could arise, let *c >* 0 denote an increase in mutant sensitivity to competition with wild types, and for simplicity assume this cost disappears in the presence of invaders. Such a cost could emerge, for instance, if wild types are more vigilant when invaders are present (Lima et al., 2021), thereby reducing the competitive impact of wild types on mutants. If mutants are rare with a positive birth rate, and wild type abundance is at equilibrium (i.e., 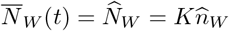), the pre-invasion mutant growth rate is 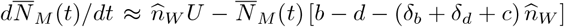. Solving 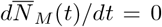 for 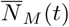 then gives a mutant equilibrium abundance of 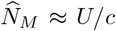, which resembles the deterministic abundance of deleterious standing variants used in other ER models (Orr & Unckless, 2008; Tomasini & Peischl, 2020). Thus, the model herein can be viewed as one in which the pre-invasion cost of invader resistance is a reduction in competitive ability equal to the expected maximum mutation rate, *U* . A natural extension of the present study would be to compare how alternative mutant fitness costs (e.g., density-independent vs. density-dependent) differentially affect *P*_*ER*_ when they act not only on initial mutant abundance but also on mutant fitness throughout ER.

Finally, the derivation of *P*_*ER*_ presented here entails its own set of assumptions. For ER via standing variation, mutants are assumed to be selectively neutral until the wild type half-life, after which wild type displacement is assumed rapid enough for mutant fitness to be positive and density-independent. For ER via *de novo* mutation, by contrast, wild type displacement is assumed slow enough for mutants to maintain a constant density-dependent growth rate throughout establishment. The overall effect is that, while the approximation remains qualitatively predictive across the explored parameter space, it loses notable quantitative accuracy under moderate invader growth rates (Figure 2). A challenge for future theory is to develop analytical approaches that better account for the time-varying nature of mutant fitness while remaining interpretable. Such approaches could help build intuition about the eco-evolutionary feedbacks underlying ER in more complex scenarios, and might also offer simple rules of thumb regarding which resident–invader interactions populations are most likely to withstand through natural selection.

## Appendix A. Analytical approximation of *P*_*ER,SV*_

For the approximation of 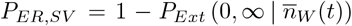 (Eq. (6)), invaders are assumed initially rare enough for 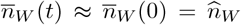 until some critical time, *t*_*c*_, at which 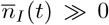. Further, invasion is assumed fast enough such that (i) 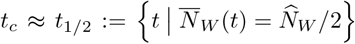, and (ii) 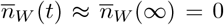 once *t > t*_1*/*2_. Under these assumptions, 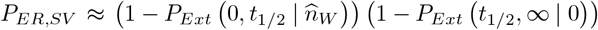, where the key quantity is

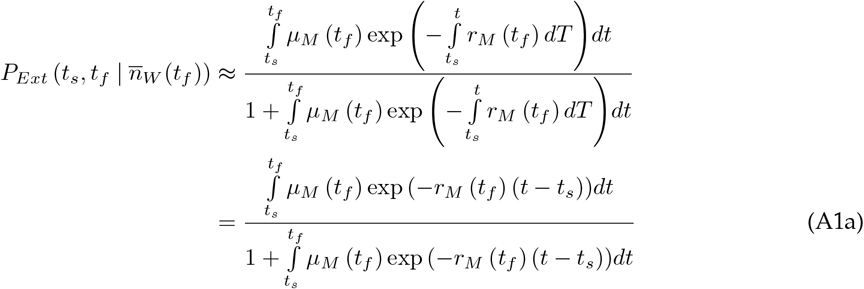

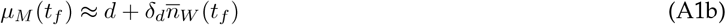

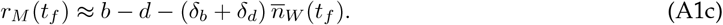

If *t*_*s*_ = 0 and 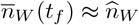 such that 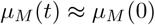 and *r*_*M*_ (*t*) ≈ *r*_*M*_ (0) = 0, Eq. (A1a) becomes

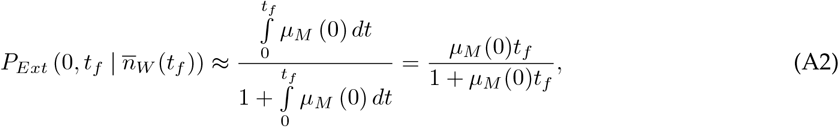

but otherwise—after substituting *z* = *t* − *t*_*s*_ and *d*_*z*_ = *dt*—Eq. (A1a) is

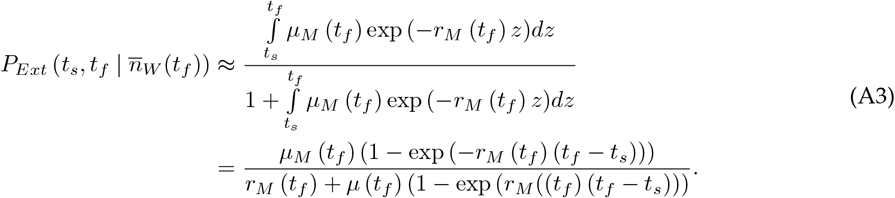

Using Eq. (8) for 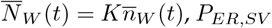 is thus

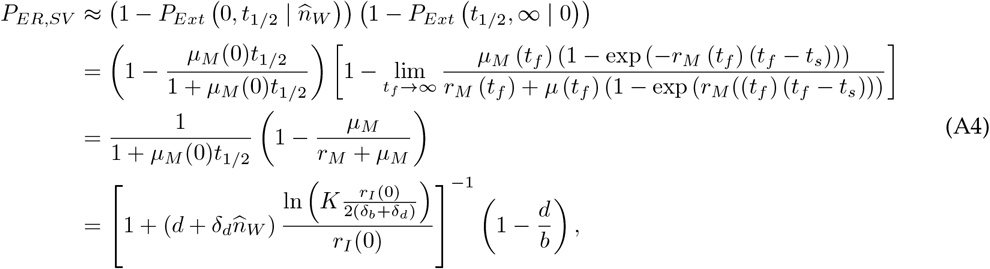

with *µ*_*M*_ = *d* and *r*_*M*_ = *b* − *d*.

## Appendix B. Analytical approximation of *P*_*ER,DN*_

For the approximation of *P*_*ER,DN*_ ≈ 1−exp (−Λ) and therefore 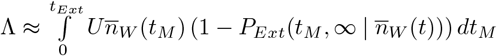, invasion is assumed slow enough for 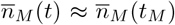 throughout mutant establishment. Equation (9) is used for 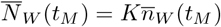 and Eq. (A3) for 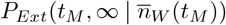, such that

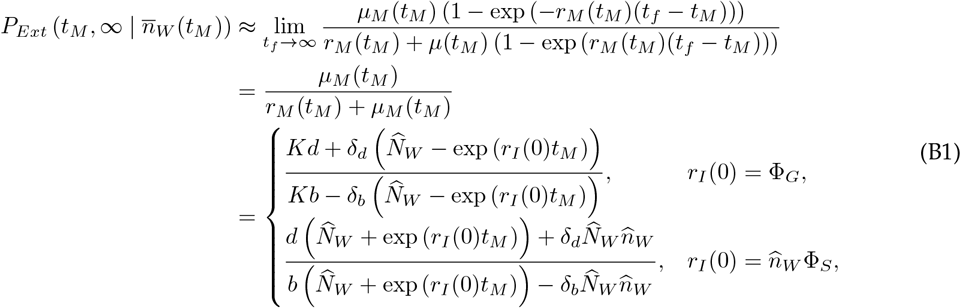

where 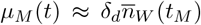 and 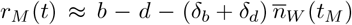. In what follows, first considered are generalist invaders with *r*_I_ (0) = Φ_G_, then specialists with 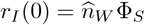.

### Generalist invaders (*r*_*I*_(0) = Φ_*G*_)

For the generalist invader case, where 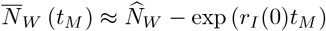 and *t*_*Ext*_ = ln 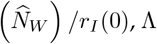 is

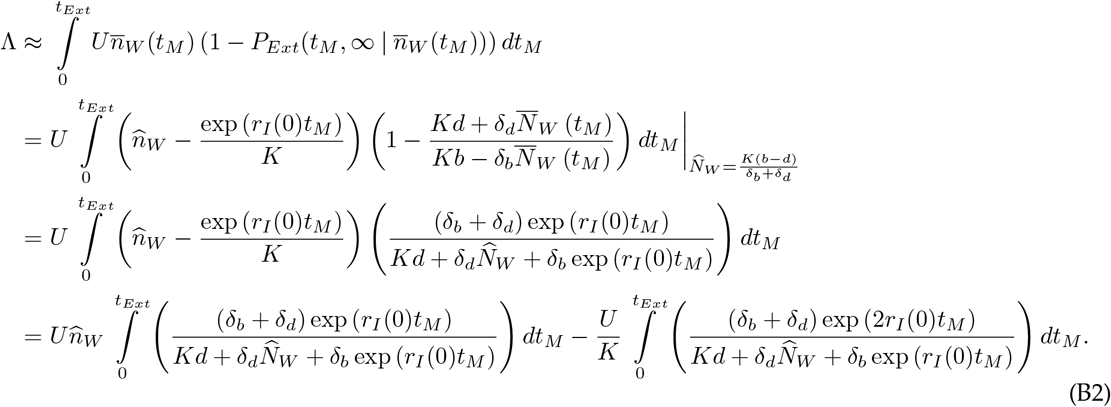

Substituting *z* = exp (*r*_*I*_ (0)*t*_*M*_) and *dz* = *zr*_*I*_ (0)*dt*_*M*_ yields

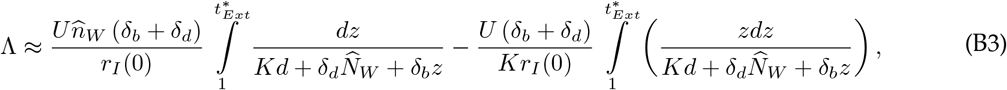

which, with 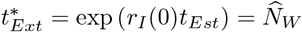, becomes

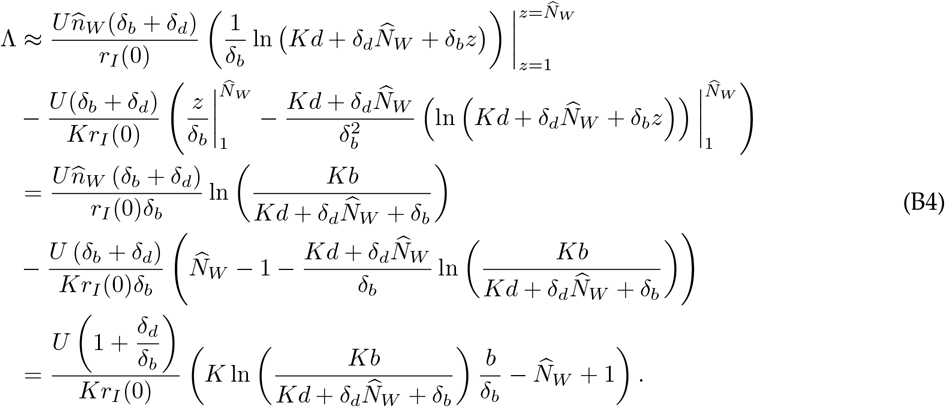

Assuming *K* ≫ 1 such that Λ is approximately equal to Eq. (B4)’s infinite-*K* limit, one obtains

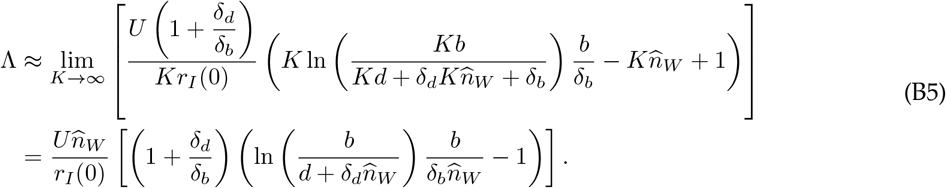

#### Specialist invaders 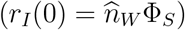

For the specialist invader case, where 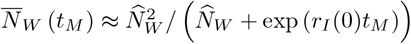 and *t*_*Ext*_ → ∞, Λ is

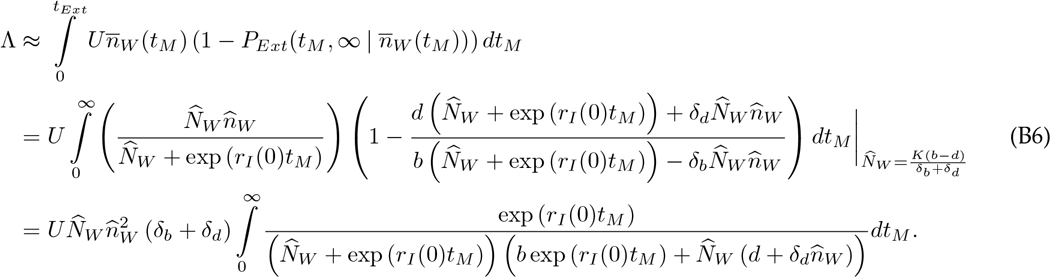

Substituting *z* = exp (*r*_*I*_ (0)*t*_*M*_) and *dz* = *zr*_*I*_ (0)*dt*_*M*_ gives

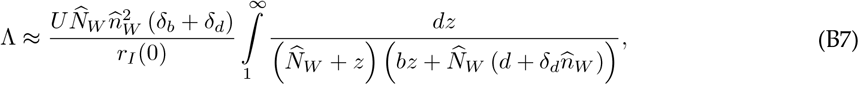

which, through partial fraction decomposition, can be written as

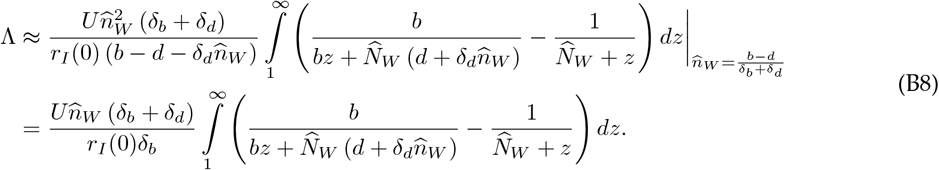

Integrating then yields

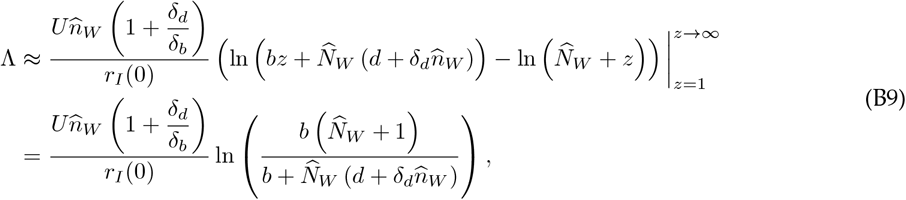

and, by assuming *K* ≫ 1, one acquires

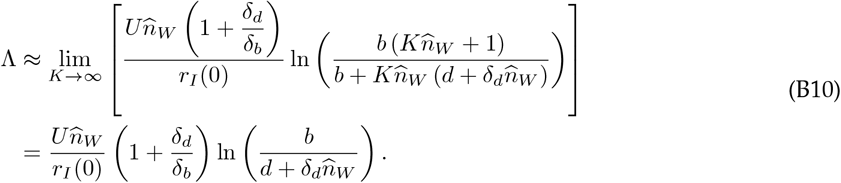

